# Germline mitotic quiescence and programmed cell death are induced in *C. elegans* by exposure to pathogenic *P. aeruginosa*

**DOI:** 10.1101/2023.08.08.552522

**Authors:** Daniel P. Bollen, Kirthi C. Reddy, Dennis H. Kim, Monica P. Colaiácovo

**Affiliations:** Division of Infectious Diseases and Department of Pediatrics, Boston Children’s Hospital and Harvard Medical School, Boston, MA 02115, USA; Department of Biology, Massachusetts Institute of Technology, Cambridge, MA 02139, USA; Department of Genetics, Harvard Medical School, Boston, MA 02115, USA

**Keywords:** germline, *C. elegans*, *P. aeruginosa*, pathogen, programmed cell death, meiosis

## Abstract

The impact of exposure to microbial pathogens on animal reproductive capacity and germline physiology is not well understood. The nematode *Caenorhabditis elegans* is a bacterivore that encounters pathogenic microbes in its natural environment. How pathogenic bacteria affect host reproductive capacity of *C. elegans* is not well understood. Here, we show that exposure of *C. elegans* hermaphrodites to the Gram-negative pathogen *Pseudomonas aeruginosa* causes a marked reduction in brood size with concomitant reduction in the number of nuclei in the germline and gonad size. We define two processes that are induced that contribute to the decrease in the number of germ cell nuclei. First, we observe that infection with *P. aeruginosa* leads to the induction of programmed germ cell death. Second, we observe that this exposure induces mitotic quiescence in the proliferative zone of the *C. elegans* gonad. Importantly, these processes appear to be reversible; when animals are removed from the presence of *P. aeruginosa*, germ cell death is abated, germ cell nuclei numbers increase, and brood sizes recover. The reversible germline dynamics during exposure to *P. aeruginosa* may represent an adaptive response to improve survival of progeny and may serve to facilitate resource allocation that promotes survival during pathogen infection.

## Introduction

Host survival and reproductive capacity during infection contribute to evolutionary fitness. How infection influences reproduction and germline physiology is not well understood. The bacterivorous nematode *Caenorhabditis elegans* encounters a wide range of pathogenic microbes in its native environment (Schulenburg and Félix 2017). *C. elegans* responds to infection through innate immune signaling (Kim and Ewbank 2018) and behavioral avoidance mechanisms (Kim and Flavell 2020).

*Pseudomonas aeruginosa* is an environmental bacterium that resides in soil and water and can cause devastating opportunistic infections in humans. *P. aeruginosa* is capable of infecting an evolutionarily diverse range of host organisms (Campa et al. 1993; Rahme et al. 1995; Reynolds and Kollef 2021). Infection of *C. elegans* by *P. aeruginosa* has been studied for over twenty years, and *P. aeruginosa* has been shown to kill *C. elegans* under a range of experimental conditions (Mahajan-Miklos et al. 1999; Tan et al. 1999). Its long history of study and the existence of tools such as an ordered transposon insertion mutant library (Liberati et al. 2006) make it an attractive and tractable model pathogen to study in this context.

We sought to understand how reproduction of *C. elegans* hermaphrodites is affected by exposure to *P. aeruginosa.* It has been reported that pathogenic bacteria such as *Shigella spp.*, *Burkholderia spp.*, *Microbacterium sp.*, *Staphylococcus aureus*, and *Serratia marcescens* reduce the number of progeny of *C. elegans*, but the underlying mechanisms are not understood (O’Quinn et al. 2001; Irazoqui et al. 2010; Kesika et al. 2015; Madhu et al. 2019; Le et al. 2022). Elucidation of these mechanisms may shed light on how animals sense and effectively utilize resources upon pathogen exposure.

Here, we study the effects of *P. aeruginosa* infection on the *C. elegans* germline. We show that exposure to *P. aeruginosa* induces a reduction in brood size in *C. elegans*. We find that exposure to *P. aeruginosa* induces programmed germ cell death, as well as mitotic quiescence in the proliferative region of the gonad. Importantly, these phenotypes are seen to constitute a dynamic process; they are reversible and contingent on exposure to the *Pseudomonas* lawn milieu. These responses may represent protective processes taken to ensure progeny viability, or the effects of shunting resources from reproduction to ensure host survival in the face of pathogenic stress.

## Materials and Methods

### C. elegans Strains

All *C. elegans* strains used in this study were maintained as previously described, with N2 Bristol worms used as the wildtype background (Brenner 1974). Animals were maintained and assays were performed at 20°C. The following strains were used in this study: MD701 (*bcIs39 [lim-7p::ced-1::GFP + lin-15(+)]*); ZD541 (*pmk-1 (km25); bcIs39*); ZD2678 (*cep-1 (gk138); bcIs39*); ZD2691 (*zip-2 (ok3730); bcIs39*).

### Bacteria Propagation

Plates for *P. aeruginosa* exposure assays were prepared as described (Meisel et al. 2014). Briefly, LB broth was inoculated with *P. aeruginosa* PA14 glycerol stock. Cultures were incubated overnight shaking at 37°C to allow growth to stationary phase. 200 µL of culture was seeded onto 3.5cm pre-warmed plates of slow-kill assay (SKA) media (Tan et al. 1999). Plates were tilted briefly to coat the entire agar surface with the bacterial culture. Plates were first incubated overnight at 37°C and then at room temperature for 24 hours. The same procedure was followed to prepare *E. coli* OP50 exposure plates for use as a control.

### Brood Size Assays

To measure the brood sizes of animals, age-synchronized hermaphrodite animals were grown to young adults (66 hours old) and then moved to bacteria of interest. For assays where the number of progeny produced during the exposure window was counted, animals were singled onto SKA plates seeded with the bacteria of interest, otherwise they were picked to a single shared plate for the duration of exposure.

At the end of the exposure period, animals were singled to nematode growth media (NGM) plates seeded with 100µL *E. coli* OP50 for recovery. Animals were moved every 24 hours to a fresh plate, and progeny on the vacated plates was counted. If animals were dead or missing at the end of the 24 hour period, the plate was omitted from analysis.

### Germline Apoptosis Assays

For germline programmed cell death assays a *ced-1::gfp* reporter strain was used (Zhou et al. 2001). Young adult animals were exposed to bacteria of interest for 6 hours. 20-25 animals were then mounted under a coverslip with 7 μl of 3mM sodium azide on a 1.5% agar pad. Germline nuclei enveloped with fluorescent CED-1 were observed in late pachytene and scored under 40x magnification using a Zeiss Axioimager Z1 microscope. Partially obscured gonad arms were not scored.

### Germline Size Reduction Assays

To assay germline size reduction, young adult animals were exposed to bacteria of interest for 24 hours. 20-25 animals were dissected, DAPI-stained and mounted as described previously (Colaiácovo et al. 2003). Images of one germline hemisphere were taken and stitched together with an ImageJ plugin (Preibisch et al. 2009). Nuclei were counted for each gonad arm sample, from the distal tip up to the end of late pachytene, omitting diplotene and diakinesis stage nuclei.

### Egg laying Assays

To stage eggs laid after exposure to bacteria, young adult animals (66 hours old) were exposed to bacteria of interest for 24 hours. After this exposure, animals were moved to NGM plates seeded with *E. coli* OP50, 5 per plate. 3 plates per condition were assayed. After one hour, animals were removed and the stage of the eggs laid was observed using a dissecting microscope. Stages were categorized according to a previously established rubric (Ringstad and Horvitz 2008).

### Fluorescence Immunostaining

Whole mount preparation of dissected gonads, fixation, and immunostainings were performed as described previously (Colaiácovo et al. 2003). An α-phospho-histone H3 (Ser10) primary rabbit antibody (Cell Signaling) was used at 1:2000 dilution. The secondary antibody used was Alexa-488 goat α-rabbit (Jackson ImmunoResearch Laboratories) at a 1:500 dilution. Vectashield from Vector Laboratories (Burlingame, CA) was used as a mounting media and anti-fading agent.

### Imaging

Images of whole-mounted gonads were collected at 0.5 µm z-intervals with an IX-70 microscope (Olympus) and a cooled CCD camera (model CH350; Roper Scientific) controlled by the DeltaVision system (Applied Precision). Images were deconvolved using the SoftWorx 3.3.6 software (Applied Precision) and processed with Fiji ImageJ (Schindelin et al. 2012).

### Statistical Analysis

Statistical tests were carried out using Graphpad Prism. Tests used are described in figure legends. Prior to statistical comparisons, outliers were detected using ROUT (Q = 1%) and removed from analysis.

## Results and Discussion

### Exposure to *P. aeruginosa* results in a reduction in *C. elegans* brood size

To quantify the effect of exposure to *P. aeruginosa* on progeny production, we exposed young adult animals to *P. aeruginosa* PA14 for 24 h. All exposures in this study were done with agar plates completely seeded with the bacterial lawn to prevent the confounding effects of pathogen avoidance. We found that for the duration of the exposure, animals on the *P. aeruginosa* lawn laid few eggs when compared to animals on the standard laboratory food source, non-pathogenic *E. coli* OP50 (Figure 1A). These data were consistent with a previous report that *C. elegans* feeding on *P. aeruginosa* produced few progeny (Tan et al. 1999).

**Figure 1.**
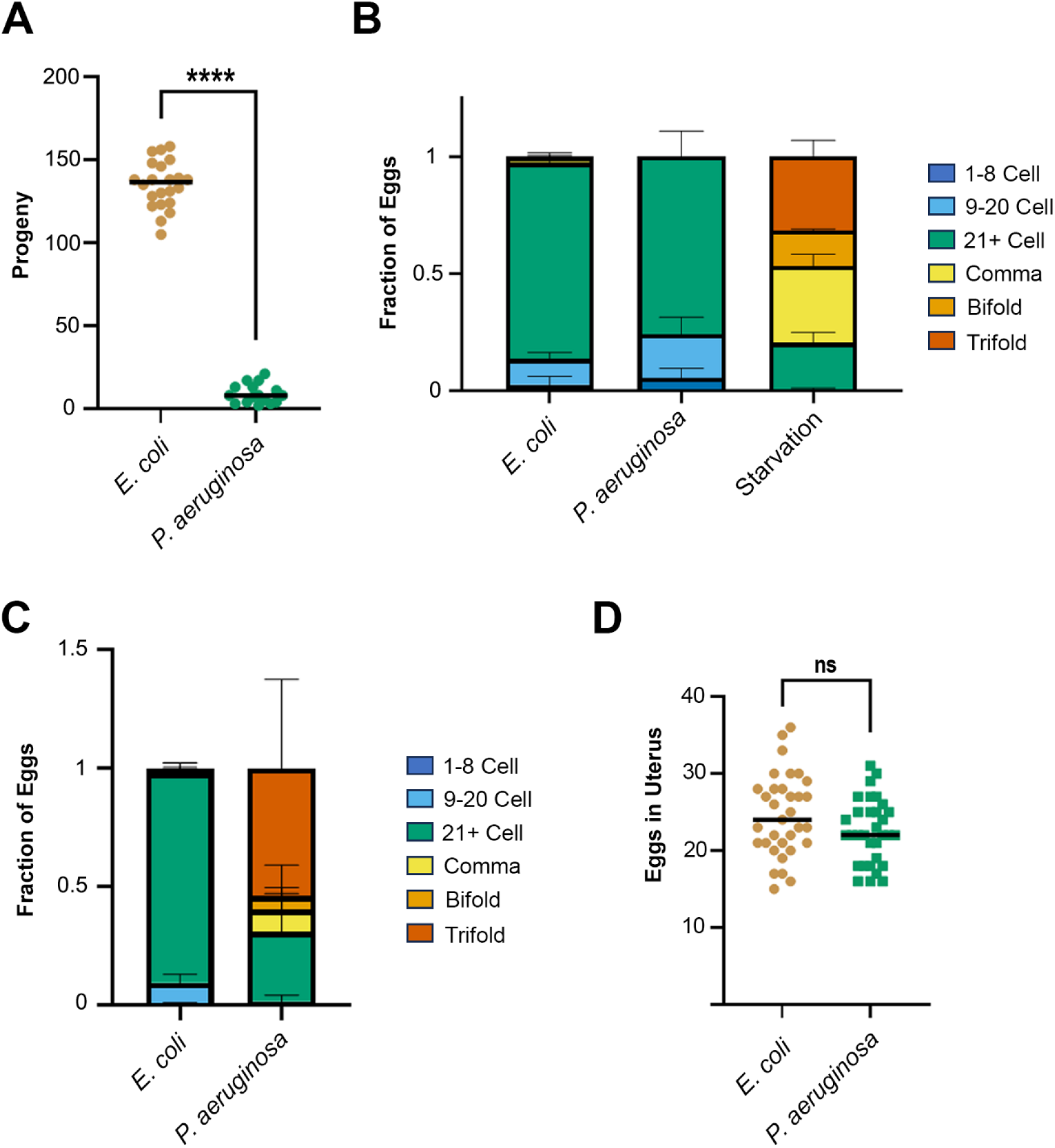
Exposure to P*. aeruginosa* results in decreased brood size that is independent of egg retention in the uterus. **A)** *C. elegans* lays significantly fewer eggs on *P. aeruginosa* compared to *E. coli*. Young adult hermaphrodites were singled onto SKA plates seeded with *E. coli* OP50 and *P. aeruginosa* PA14 for 24 hours. At the end of the exposure time the number of eggs laid were counted. Each point represents a unique animal (n ≈ 20). Welch’s t-test, two tailed, **** *p* < 0.0001. **B)** *C. elegans* does not retain eggs on *P. aeruginosa* in short timescales. Young adult hermaphrodites were exposed to *P. aeruginosa*, *E. coli*, or were starved for 6 hours. Animals were then moved to *E. coli* OP50 to lay eggs for 1 hour. Animals exposed to *P. aeruginosa* did not lay eggs at later stages compared to *E. coli*, though starved animals did. Average of three experiments with at least 50 eggs scored per condition per experiment from 15 animals for each condition. **C)** Animals exposed to *P. aeruginosa* do retain eggs after longer exposure times. Young adult hermaphrodites were exposed to *P. aeruginosa* or *E. coli* for 24 hours. Animals were then moved to *E. coli* OP50 to lay eggs for 1 hour. Animals exposed to *P. aeruginosa* laid eggs at later stages. Average of three experiments with at least 25 eggs scored per condition per experiment from 25 animals for each condition. **D)** Egg laying defect in response to *P. aeruginosa* does not result in a significant increase of eggs in the uterus. Young adult hermaphrodites were exposed to *P. aeruginosa* or *E. coli* for 24 hours. Animals were anesthetized and the number of eggs in the uterus of each animal was counted. Each point represents a unique animal (n ≈ 30). Student’s t-test, two-tailed, ns = not significant.

We investigated whether defective egg laying and egg retention was induced by exposure to *P. aeruginosa*. In hermaphrodite animals with defects in egg-laying, eggs are retained in the uterus and laid at later developmental stages (Trent et al. 1983). As such, egg retention can be measured by identifying the developmental stage of recently laid eggs (Ringstad and Horvitz 2008). To determine if the reduction in brood size was strictly due to a retention of eggs in the uterus, we exposed animals to *P. aeruginosa*, then assayed the developmental stage of eggs laid. We found that compared to the canonical egg retention induced by removal of food (Trent et al. 1983), exposure to *P. aeruginosa* caused no egg-laying deficit after 6 h (Figure 1B). However, after a longer exposure time of 24 h, a remarkable egg-laying deficit appeared when compared to what we observed for hermaphrodites exposed to *E. coli* OP50 (Figure 1C). We also noted that the number of eggs in the uterus did not increase after exposure to *P. aeruginosa*, suggesting that the missing progeny were not simply retained, and that fewer eggs were produced overall (Figure 1D). Whereas these data suggest that exposure to *P. aeruginosa* causes a defect in egg laying that may contribute to decreased progeny production, our observations also suggest that there is a reduction in the number of fertilized oocytes during exposure to *P. aeruginosa*.

### Exposure to *P. aeruginosa* induces germline shrinking

We next assessed whether the reduction in progeny upon exposure to *P. aeruginosa* might be due to physiological changes in the germline. The *C. elegans* reproductive system consists of two gonad arms connected to a uterus (Figure 2A). Nuclei are positioned in a spatiotemporal gradient along each gonad arm with a stem cell niche at the distal end giving rise to a population of mitotically proliferating nuclei (the pre-meiotic tip or proliferative zone) which then progress out of mitosis into meiosis (Kimble and Crittenden 2005 Aug 15).

**Figure 2.**
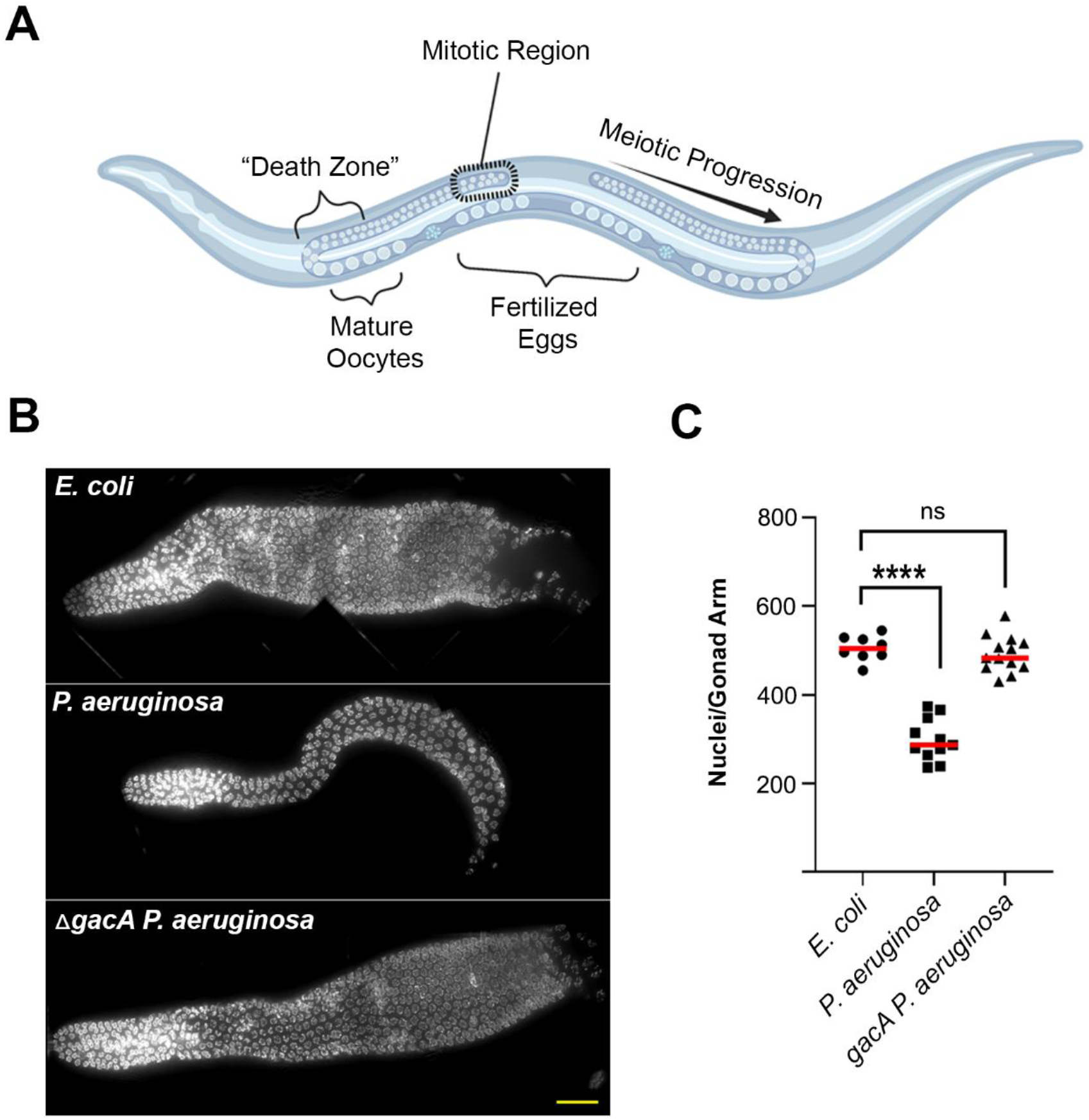
C. elegans germline undergoes shrinkage upon exposure to *P. aeruginosa.* **A)** Diagram of *C. elegans* reproductive system, with its two symmetrical gonad arms. The mitotic region is highlighted, as well as the “death zone” where nuclei can undergo apoptosis. Germline nuclei that do not undergo apoptosis complete oogenesis into mature oocytes and are fertilized, ultimately passing into the uterus where they will develop until laid. Diagram created with BioRender.com. **B)** Germlines of animals exposed to *P. aeruginosa* are reduced in size compared to those of animals exposed to *E. coli* and non-pathogenic *P. aeruginosa* mutant *gacA*. DAPI-stained germlines of animals exposed to bacteria of interest for 24 hours. Scale bar, 25µm. **C)** Quantification of the number of germline nuclei after 24 hour exposure to bacteria of interest. There is a significant reduction for animals exposed to *P. aeruginosa*, but not those exposed to the non-pathogenic *P. aeruginosa* mutant *gacA.* Welch’s t test, **** p < 0.0001, ns = not significant (n ≈ 10).

A dramatic reduction in the size of the gonads was observed for animals exposed to pathogen (Figure 2B). Notably, this reduction was absent in animals exposed to the *P. aeruginosa gacA* mutant that is defective in virulence (Rahme et al. 1995). This decrease in size could be due to a reduction in either the cytoplasmic material in the gonadal syncytium or in the number of germline nuclei. To identify changes in the number of nuclei in the germline, animals exposed to *P. aeruginosa* PA14 for 24 h were dissected, and germlines were isolated and DAPI-stained. Quantification of the number of DAPI-stained nuclei in the germline revealed a significant reduction in the number of nuclei in the gonad arms upon exposure to pathogenic *P. aeruginosa*, but notably no significant decrease in animals exposed to the *P. aeruginosa gacA* mutant (Figure 2C). This suggests that the pathogenicity of *P. aeruginosa* is responsible for the decrease in germline nuclei, as opposed to a difference in bacterial food quality between *E. coli* OP50 and *P. aeruginosa* PA14.

### Induction of germline programmed cell death in response to *P. aeruginosa* is linked to increased pathogen susceptibility but not activation of a p53/CEP-1-dependent DNA damage response pathway

Possible explanations for this reduction in germline nuclei include increased oocyte maturation rate, a reduction in mitotic proliferation, or an increase in germline apoptosis rates. Due to our observation of a reduction in the number of eggs produced during exposure, we considered an increased oocyte maturation rate unlikely. To determine if germline apoptosis plays a role in the reduction in germline size, we measured rates of germline programmed cell death (PCD) using a *ced-1::gfp* reporter strain. *ced-1* encodes for a phagocytic corpse-recognition engulfment protein essential in the apoptotic process. Nuclei engulfed during programmed cell death become enveloped in CED-1 (Figure 3A) (Zhou et al. 2001). There is a basal physiological rate of germline PCD, and this can be increased upon exposure to chemical, osmotic, or heat stress, as well as starvation (Gumienny et al. 1999; Salinas et al. 2006). We found that after 6 h of exposure to *P. aeruginosa* PA14, a greater number of CED-1-positive nuclei appear compared to *E. coli* OP50 (Figure 3B). Importantly, *ΔgacA* PA14 did not induce germline PCD, again suggesting that these effects are due to pathogenic infection, in line with our observation for gonad size reduction. We found similar results for animals exposed to *P. aeruginosa* mutants defective in other virulence factors, such as *rhlR and lasR* (Figure 3B) (Kariminik et al. 2017). Exposure of *C. elegans* to *P. aeruginosa* PA14 was previously reported to not cause increased germline PCD (Aballay and Ausubel 2001). The discrepancy with our results may be due to technical issues related to interactions between the *P. aeruginosa* PA14 lawn and the dye used to stain for corpses, which was alluded to in the prior study (Aballay and Ausubel 2001).

**Figure 3.**
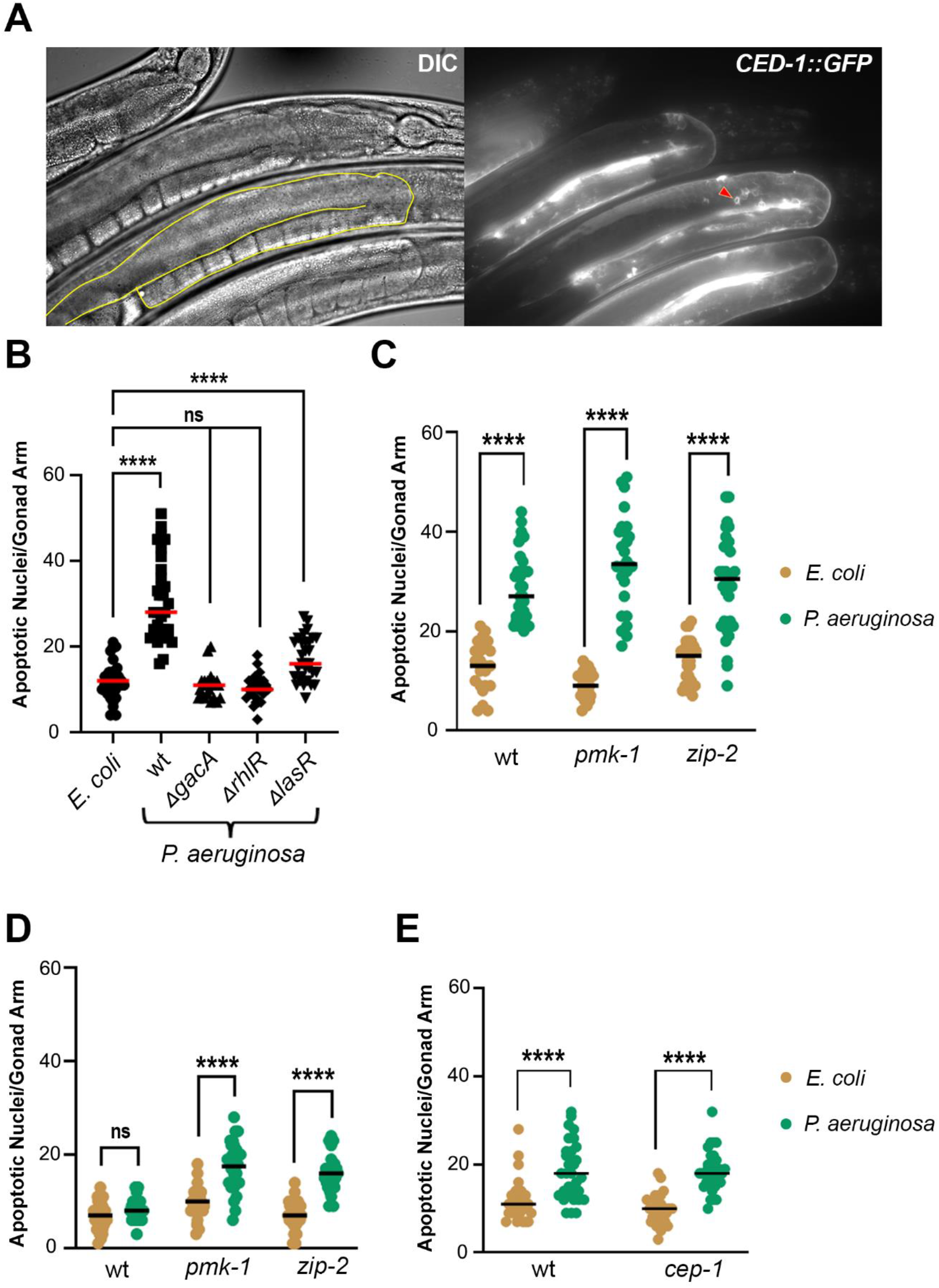
Exposure to *P. aeruginosa* induces apoptosis in the *C. elegans* germline. **A)** Representative image of *ced-1::gfp* reporter strain MD701 on *E. coli*. Germline outline in yellow on DIC. Apoptotic nuclei can be seen in the death zone of the germline enveloped with CED-1::gfp, an example of which is marked with a red arrowhead. **B)** Quantification of apoptotic germ cell nuclei after 6 h exposure to bacteria of interest. Animals exposed to *P. aeruginosa* showed induction of germline PCD which was not observed in mutants in pathogenic regulators *gacA*, *rhlR*, and *lasR*. **C)** Mutants in infection response *pmk-1* and *zip-2* show a greater magnitude of induction of germline PCD upon exposure to *P. aeruginosa*. Animals were exposed to bacteria of interest for 6 hours. **D)** Mutants in infection response *pmk-1* and *zip-2* show more rapid induction of PCD upon exposure to *P. aeruginosa*. Animals were exposed to bacteria of interest for 1 hour. **E)** p53 homolog CEP-1 is not necessary for induction of germline PCD upon exposure to *P. aeruginosa*. Welch’s t-test used for all statistical comparisons, **** p < 0.0001, *** p < 0.001, ns = not significant (n ≈ 30 gonads per condition).

Germline PCD in response to infection by *Salmonella enterica* has been reported to be dependent on PMK-1 p38 mitogen-activated protein kinase (MAPK) (Aballay et al. 2003). In contrast, we observed that upon exposure to *P. aeruginosa,* PMK-1 was not required for induction of germline PCD (Figure 3C). Moreover, animals defective in *pmk-1* showed a greater induction of germline PCD than wildtype animals. Because *pmk-1* mutants are more susceptible to pathogen infection (Kim et al. 2002), we hypothesized that the greater induction of germline PCD upon exposure to *P. aeruginosa* could be due to increased susceptibility to pathogenesis. To test this, we examined germline PCD in a *zip-2* mutant. The bZIP transcription factor ZIP-2 regulates the innate immune response of *C. elegans* independently of *pmk-1* (Estes et al. 2010). Mutants in *zip-2* showed a greater induction of germline PCD than wild-type animals after a 6 h exposure, similar to what was observed for *pmk-1* mutants, suggesting that the increased pathogen susceptibility may be associated with the severity of the germline PCD induction. To further explore this effect, we examined induction of germline PCD at earlier time points. We found that even within 1 h of exposure, both *zip-2* and *pmk-1* mutant animals showed induction of germline PCD in response to *P. aeruginosa*, while wild-type animals did not (Figure 3D), indicative of accelerated kinetics of induction of germline PCD in mutants defective in the innate immune response. The differences in our observations here and prior reports of PMK-1-dependent germline PCD during *Salmonella* infection (Aballay et al. 2003) may be due to differences in how each pathogen induces germline PCD or differences in experimental conditions.

One possibility for this induction of germline PCD is increased DNA damage among meiotic nuclei in the germline. In *C. elegans*, the p53 homolog CEP-1 is necessary for induction of germline PCD in response to DNA double-strand breaks (DSBs) caused by ionizing radiation and in mutants with either elevated programmed meiotic DSBs or impaired DSB repair (Schumacher et al. 2001). Our data shows that *cep-1* mutant animals exhibited no defect in the induction of germline PCD in response to *P. aeruginosa* (Figure 3E). This suggests that the increased germline PCD following exposure to *P. aeruginosa* is not in response to DNA damage and must occur by some other mechanism.

### Exposure to *P. aeruginosa* induces mitotic quiescence in the proliferative zone of the germline

Another possible process by which the number of germline nuclei may be reduced is a cessation of mitosis in the proliferative zone of the gonad arm. Starvation of *C. elegans* was previously observed to result in the induction of mitotic quiescence, as detected by a loss of nuclei positive for histone H3 phosphorylation, a marker indicative of M phase (Seidel and Kimble 2015).

To determine if mitotic quiescence was occurring in response to *P. aeruginosa* exposure, young adult animals were exposed to *P. aeruginosa* PA14 for 6 h, and their whole mounted gonads immunostained for phospho-histone H3. We found that phospho-histone H3-positive nuclei were almost completely absent from *P. aeruginosa*-exposed animals, though not in animals exposed to *E. coli* OP50 or *ΔgacA P. aeruginosa* PA14 (Figure 4A, B). This suggests that exposure to pathogen induces mitotic quiescence in *C. elegans*.

**Figure 4.**
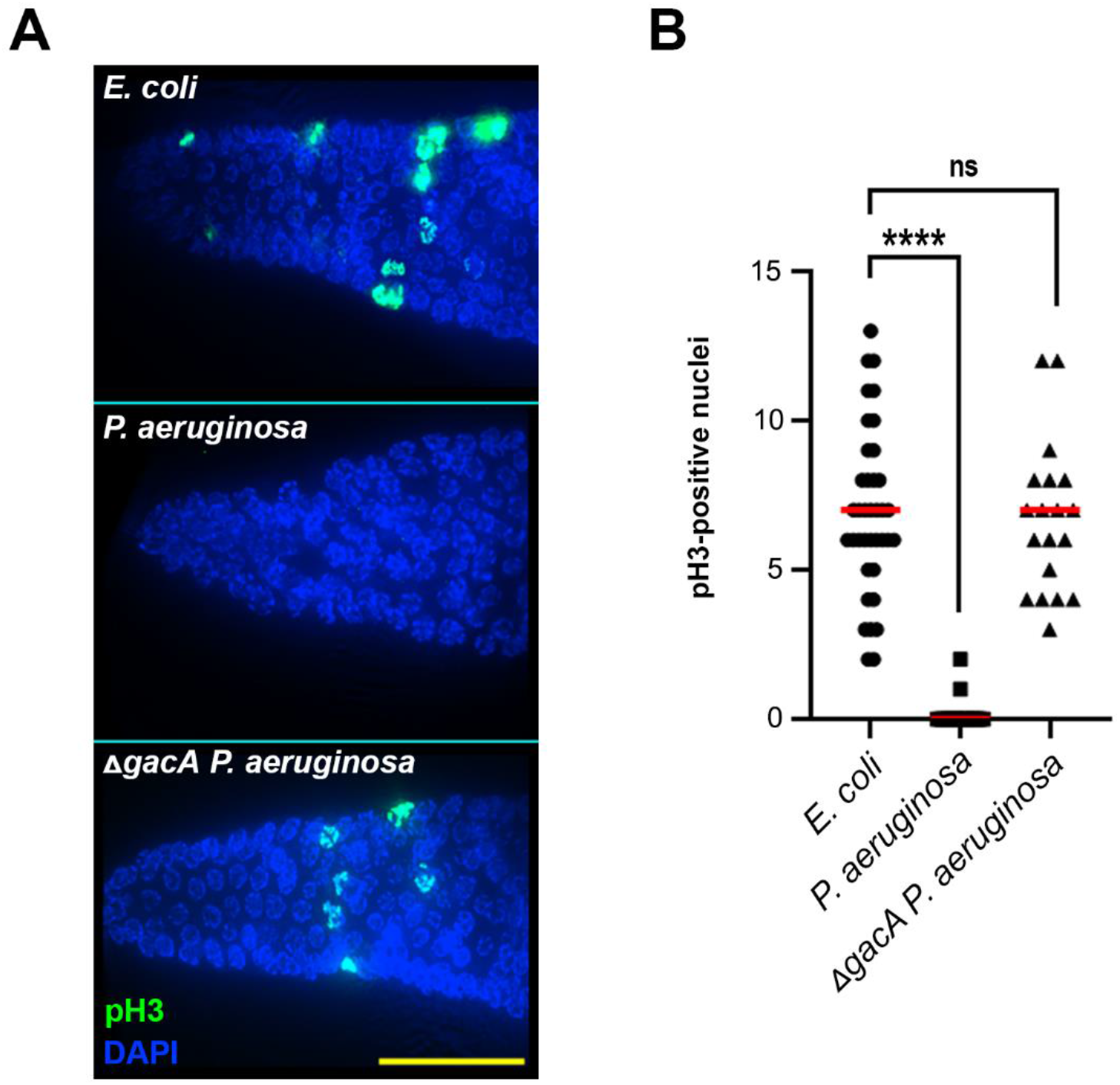
Exposure to *P. aeruginosa* induces mitotic quiescence in the *C. elegans* germline. **A)** Representative high-resolution images of nuclei in the pre-meiotic region of the germline immunostained for phospho-histone H3 (Ser10) (pH3). Scale bar, 25 µm. **B)** Quantification of pH3 immunostaining shows loss of mitotically dividing nuclei upon exposure to *P. aeruginosa* but not in the non-pathogenic *gacA* mutant. Mann-Whitney test, **** p < 0.0001, ns = not significant (n ≈ 20 gonads scored for each).

### Pathogen-induced germline changes are reversible upon cessation of exposure to *P. aeruginosa*

To determine if the changes in the germline that we observed during exposure to *P. aeruginosa* were reversible, we exposed animals to *P. aeruginosa* PA14 and then returned them to *E. coli* OP50. After 24 h of exposure to *P. aeruginosa*, while animals showed reduced egg-laying capacity during the next 24 h period, animals transferred back to *E. coli* OP50 recovered and ultimately laid a similar number of eggs to animals exposed only to *E. coli* OP50 (Figure 5A). In addition, animals transferred from *P. aeruginosa* to *E. coli* OP50 exhibited phospho-histone H3-positive nuclei within 2 h (Figure 5B). This two-hour time scale is consistent with the recovery time sufficient for recovery from starvation-induced mitotic quiescence (Seidel and Kimble 2015). We also observed that a 2 h recovery on *E. coli* OP50 after transfer from *P. aeruginosa* was sufficient to reduce the number of apoptotic nuclei in the gonad arms (Figure 5C). The relatively rapid recovery in the levels of germline PCD induction suggests that exposure to the pathogenic lawn, and not necessarily the longer-term effects of damage from infection and a subsequent innate immune response, contributes to the induction of germline PCD. We further observed that animals removed from *P. aeruginosa* and placed on *E. coli* OP50 exhibited recovery of the germline, with nuclei numbers increasing over several days (Figure 5D).

**Figure 5.**
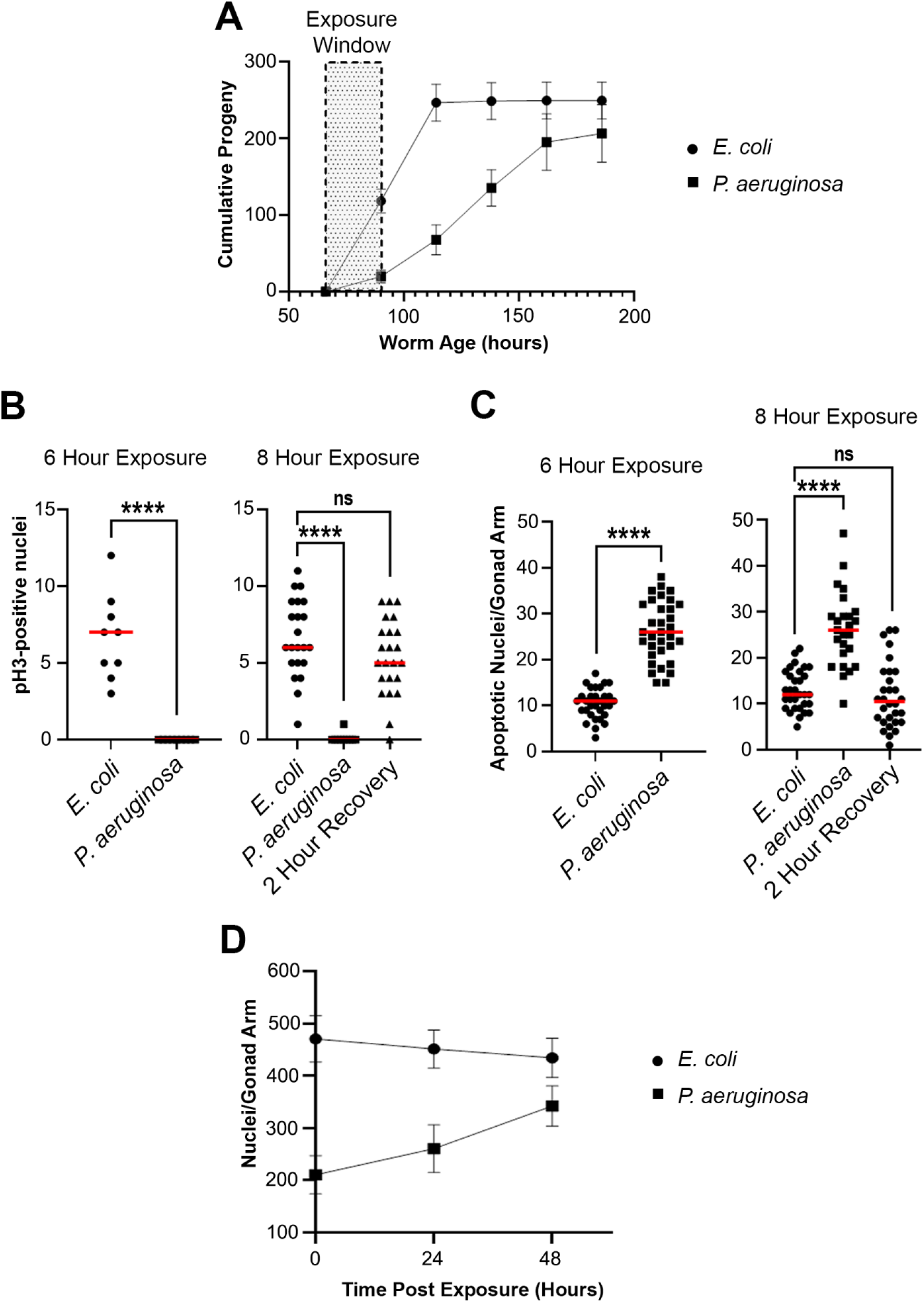
Germline changes in response to *P. aeruginosa* are reversible upon removal of *P. aeruginosa*. **A)** Young adult animals exposed to *P. aeruginosa* show sustained but reversible reduction in brood size. Animals were exposed to bacteria of interest for 24 hours, then moved to *E. coli* NGM plates to recover. Animals were moved every 24 hours to a fresh plate and the number of progeny produced was counted. Points represent the average of 25 animals. Error bars represent the standard deviation. **B)** Mitotic quiescence is abated within 2 hours of recovery on *E. coli*. Animals were exposed to bacteria of interest for 6 hours. A subset of animals exposed to *P. aeruginosa* were allowed to recover on *E. coli* for 2 hours, then phospho-histone H3-positive nuclei were assayed again. Mann-Whitney test was used for non-parametric comparisons, and student’s t-test was used to compare parametric data, **** p < 0.0001, ns = not significant (n ≈ 15 gonads per condition). **C)** Germline PCD induction is abated within 2 hours of recovery on *E. coli*. *ced-1::gfp*, animals were exposed to bacteria of interest for 6 h, assessed for the number of apoptotic nuclei per gonad arm, and then assessed again 2 hours later, with a subset of animals exposed to *P. aeruginosa* allowed to recover on *E. coli* for 2 hours. **** p < 0.0001, Welch’s t-test (n ≈ 30 gonads per condition). **D)** Gonad size recovers in animals removed from *P. aeruginosa*. Animals were exposed to bacteria of interest for 24 hours and then the number of DAPI-stained nuclei in each dissected gonad was scored. Animals were then moved to *E. coli* OP50 to recover and were dissected 24 h and 48 hours later. Each point represents the average of n ≈ 15 animals. Error bars represent the standard deviation.

These observations show that exposure to pathogenic bacteria results in a reduced number of progeny laid, with a concomitant increase in germline programmed cell death and induction of mitotic quiescence in the germline. These observations are accompanied by a reduction in the number of nuclei in the germline, and these phenomena appear to be reversible once the animals are in the presence of non-pathogenic bacteria. We speculate that the combination of germline PCD and mitotic quiescence in response to exposure to *P. aeruginosa* may represent a protective program to maintain brood viability in the presence of a pathogenic microbial environment, with a reduction in the number of progeny laid in an unfavorable microbial environment to preserve progeny viability and promote robust brood sizes when there has been a transition to a more beneficial microbial environment.

This suite of phenotypes is also known to occur in response to starvation (Salinas et al. 2006; Angelo and Gilst 2009; Seidel and Kimble 2015). Furthermore, reports have shown that exposure to *P. aeruginosa* reduces lipid stores in *C. elegans* (Nhan et al. 2019) and leads to downregulation of the acyl-CoA dehydrogenase *acdh-1* (Fletcher et al. 2019) which has previously been identified as a marker of the starvation response (Van Gilst et al. 2005). As such, these phenotypes bare some resemblance to adult reproductive diapause (ARD). However, the lethal nature of *P. aeruginosa* exposure does not allow for the same long-term reproductive quiescence reported in ARD (Angelo and Gilst 2009; Seidel and Kimble 2011; Gerisch et al. 2020). While our experiments with non-pathogenic *P. aeruginosa* mutants shows that the causative factors in these changes are not due to an underlying nutritive difference between two bacterial species, the similarities in the germline phenotypes induced by these distinct stressors suggest that there may be commonalities in underlying host responses to trigger reversible germline changes. Recent research into immunometabolic crosstalk has uncovered relationships between metabolic stress and immune activation (Ayres 2019; Penkov et al. 2019; Anderson and Pukkila-Worley 2020; Peterson et al. 2022). The similarities between adult reproductive diapause and our here described germline pathogen response may arise from this immunometabolic crosstalk.

While here we describe a set of reversible germline phenotypes accompanying a reduction in progeny number upon exposure to *P. aeruginosa*, the molecular factors involved remain elusive. Further study of the pathogen and host-associated factors necessary for this germline response may yield insight into how animals sense and effectively respond to pathogens.

## Data Availability

Strains and reagents are available upon request. The authors affirm that all data necessary for confirming the conclusions of the article are present within the article, figures, and tables.

## Acknowledgments

Some strains were provided by the Caenorhabditis Genetics Center, which is funded by NIH Office of Research Infrastructure Programs (P40 OD010440). We also thank members of the Kim and Colaiácovo labs for helpful conversations and feedback on this work.

## References

Aballay A, Ausubel FM. 2001. Programmed cell death mediated by ced-3 and ced-4 protects Caenorhabditis elegans from Salmonella typhimurium-mediated killing. Proc Natl Acad Sci U S A. 98(5):2735–2739. doi:10.1073/pnas.041613098.

Aballay A, Drenkard E, Hilbun LR, Ausubel FM. 2003. Caenorhabditis elegans Innate Immune Response Triggered by Salmonella enterica Requires Intact LPS and Is Mediated by a MAPK Signaling Pathway. Current Biology. 13(1):47–52. doi:10.1016/S0960-9822(02)01396-9.

Anderson SM, Pukkila-Worley R. 2020. Immunometabolism in Caenorhabditis elegans. PLoS Pathog. 16(10):e1008897. doi:10.1371/journal.ppat.1008897.

Angelo G, Gilst MR Van. 2009. Starvation Protects Germline Stem Cells and Extends Reproductive Longevity in C. elegans. Science (1979). 326(5955):954–958. doi:10.1126/science.1178343.

Ayres JS. 2019. Immunometabolism of infections. Nature Reviews Immunology 2019 20:2. 20(2):79–80. doi:10.1038/s41577-019-0266-9.

Brenner S. 1974. The genetics of Caenorhabditis elegans. Genetics. 77(1):71–94. doi:10.1093/genetics/77.1.71.

Campa M, Bendinelli M, Friedman H, editors. 1993. Pseudomonas aeruginosa as an Opportunistic Pathogen. Boston, MA: Springer US.

Colaiácovo MP, MacQueen AJ, Martinez-Perez E, McDonald K, Adamo A, La Volpe A, Villeneuve AM. 2003. Synaptonemal Complex Assembly in C. elegans Is Dispensable for Loading Strand-Exchange Proteins but Critical for Proper Completion of Recombination. Dev Cell. 5(3):463–474. doi:10.1016/S1534-5807(03)00232-6.

Estes KA, Dunbar TL, Powell JR, Ausubel FM, Troemel ER. 2010. bZIP transcription factor zip-2 mediates an early response to Pseudomonas aeruginosa infection in Caenorhabditis elegans. Proceedings of the National Academy of Sciences. 107(5):2153–2158. doi:10.1073/pnas.0914643107.

Fletcher M, Tillman EJ, Butty VL, Levine SS, Kim DH. 2019. Global transcriptional regulation of innate immunity by ATF-7 in C. elegans. Garsin DA, editor. PLoS Genet. 15(2):e1007830. doi:10.1371/journal.pgen.1007830.

Gerisch B, Tharyan RG, Mak J, Denzel SI, Popkes-van Oepen T, Henn N, Antebi A. 2020. HLH-30/TFEB Is a Master Regulator of Reproductive Quiescence. Dev Cell. 53(3):316–329.e5. doi:10.1016/j.devcel.2020.03.014.

Van Gilst MR, Hadjivassiliou H, Yamamoto KR. 2005. A Caenorhabditis elegans nutrient response system partially dependent on nuclear receptor NHR-49. Proc Natl Acad Sci U S A. 102(38):13496– 13501. doi:10.1073/PNAS.0506234102.

Gumienny TL, Lambie E, Hartwieg E, Horvitz HR, Hengartner MO. 1999. Genetic control of programmed cell death in the Caenorhabditis elegans hermaphrodite germline. Development. 126(5):1011–1022. doi:10.1242/dev.126.5.1011.

Irazoqui JE, Troemel ER, Feinbaum RL, Luhachack LG, Cezairliyan BO, Ausubel FM. 2010. Distinct pathogenesis and host responses during infection of C. elegans by P. aeruginosa and S. aureus. PLoS Pathog. 6(7):1–24. doi:10.1371/journal.ppat.1000982.

Kariminik A, Baseri-Salehi M, Kheirkhah B. 2017. Pseudomonas aeruginosa quorum sensing modulates immune responses: An updated review article. Immunol Lett. 190:1–6. doi:10.1016/j.imlet.2017.07.002.

Kesika P, Prasanth MI, Balamurugan K. 2015. Modulation of Caenorhabditis elegans immune response and modification of Shigella endotoxin upon interaction. J Basic Microbiol. 55(4):432–450. doi:10.1002/JOBM.201400511.

Kim DH, Ewbank JJ. 2018. Signaling in the innate immune response. WormBook.:1–35. doi:10.1895/wormbook.1.83.2.

Kim DH, Feinbaum R, Alloing G, Emerson FE, Garsin DA, Inoue H, Tanaka-Hino M, Hisamoto N, Matsumoto K, Tan M-W, et al. 2002. A conserved p38 MAP kinase pathway in Caenorhabditis elegans innate immunity. Science. 297(5581):623–6. doi:10.1126/science.1073759.

Kim DH, Flavell SW. 2020. Host-microbe interactions and the behavior of Caenorhabditis elegans. J Neurogenet. 34(3–4):500–509. doi:10.1080/01677063.2020.1802724.

Kimble J, Crittenden SL. 2005 Aug 15. Germline proliferation and its control. Strome S, editor. WormBook. doi:10.1895/wormbook.1.13.1.

Le TS, Nguyen THG, Ha BH, Huong BTM, Nguyen TTH, Vu KD, Ho TC, Wang J. 2022. Reproductive Span of Caenorhabditis Elegans is Extended by Microbacterium Sp. J Nematol. 54(1). doi:10.2478/jofnem-2022-0010.

Liberati NT, Urbach JM, Miyata S, Lee DG, Drenkard E, Wu G, Villanueva J, Wei T, Ausubel FM. 2006. An ordered, nonredundant library of Pseudomonas aeruginosa strain PA14 transposon insertion mutants. Proc Natl Acad Sci U S A. 103(8):2833–2838. doi:10.1073/pnas.0511100103.

Madhu BJ, Salazar AE, Gumienny TL. 2019. Caenorhabditis elegans egg-laying and brood-size changes upon exposure to Serratia marcescens and Staphylococcus epidermidis are independent of DBL-1 signaling. MicroPubl Biol. 2019:10.17912/2r51-b476. doi:10.17912/2R51-B476.

Mahajan-Miklos S, Tan MW, Rahme LG, Ausubel FM. 1999. Molecular Mechanisms of Bacterial Virulence Elucidated Using a Pseudomonas aeruginosa– Caenorhabditis elegans Pathogenesis Model. Cell. 96(1):47–56. doi:10.1016/S0092-8674(00)80958-7.

Meisel JD, Panda O, Mahanti P, Schroeder FC, Kim DH. 2014. Chemosensation of bacterial secondary metabolites modulates neuroendocrine signaling and behavior of C. elegans. Cell. 159(2):267–80. doi:10.1016/j.cell.2014.09.011.

Nhan JD, Turner CD, Anderson SM, Yen CA, Dalton HM, Cheesman HK, Ruter DL, Naresh NU, Haynes CM, Soukas AA, et al. 2019. Redirection of SKN-1 abates the negative metabolic outcomes of a perceived pathogen infection. Proc Natl Acad Sci U S A. 116(44):22322–22330. doi:10.1073/PNAS.1909666116.

O’Quinn AL, Wiegand EM, Jeddeloh JA. 2001. Burkholderia pseudomallei kills the nematode Caenorhabditis elegans using an endotoxin-mediated paralysis. Cell Microbiol. 3(6):381–393. doi:10.1046/J.1462-5822.2001.00118.X.

Penkov S, Mitroulis I, Hajishengallis G, Chavakis T. 2019. Immunometabolic Crosstalk: An Ancestral Principle of Trained Immunity? Trends Immunol. 40(1):1–11. doi:10.1016/J.IT.2018.11.002.

Peterson ND, Icso JD, Salisbury JE, Rodríguez T, Thompson PR, Pukkila-Worley R. 2022. Pathogen infection and cholesterol deficiency activate the C. elegans p38 immune pathway through a TIR-1/SARM1 phase transition. Elife. 11. doi:10.7554/elife.74206.

Preibisch S, Saalfeld S, Tomancak P. 2009. Globally optimal stitching of tiled 3D microscopic image acquisitions. Bioinformatics. 25(11):1463. doi:10.1093/bioinformatics/BTP184.

Rahme LG, Stevens EJ, Wolfort SF, Shao J, Tompkins RG, Ausubel FM. 1995. Common virulence factors for bacterial pathogenicity in plants and animals. Science. 268(5219):1899–1902. doi:10.1126/science.7604262.

Reynolds D, Kollef M. 2021. The Epidemiology and Pathogenesis and Treatment of Pseudomonas aeruginosa Infections: An Update. Drugs 2021 81:18. 81(18):2117–2131. doi:10.1007/S40265-021-01635-6.

Ringstad N, Horvitz HR. 2008. FMRFamide neuropeptides and acetylcholine synergistically inhibit egg-laying by C. elegans. Nature Neuroscience 2008 11:10. 11(10):1168–1176. doi:10.1038/nn.2186.

Salinas LS, Maldonado E, Navarro RE. 2006. Stress-induced germ cell apoptosis by a p53 independent pathway in Caenorhabditis elegans. Cell Death & Differentiation 2006 13:12. 13(12):2129–2139. doi:10.1038/sj.cdd.4401976.

Schindelin J, Arganda-Carreras I, Frise E, Kaynig V, Longair M, Pietzsch T, Preibisch S, Rueden C, Saalfeld S, Schmid B, et al. 2012. Fiji: an open-source platform for biological-image analysis. Nature Methods 2012 9:7. 9(7):676–682. doi:10.1038/nmeth.2019.

Schulenburg H, Félix MA. 2017. The natural biotic environment of Caenorhabditis elegans. Genetics. 206(1):55–86. doi:10.1534/genetics.116.195511.

Schumacher B, Hofmann K, Boulton S, Gartner A. 2001. The C. elegans homolog of the p53 tumor suppressor is required for DNA damage-induced apoptosis. Current Biology. 11(21):1722–1727. doi:10.1016/S0960-9822(01)00534-6.

Seidel HS, Kimble J. 2011. The Oogenic Germline Starvation Response in C. elegans. PLoS One. 6(12):e28074. doi:10.1371/journal.pone.0028074.

Seidel HS, Kimble J. 2015. Cell-cycle quiescence maintains Caenorhabditis elegans germline stem cells independent of GLP-1/Notch. Elife. 4. doi:10.7554/elife.10832.

Tan MW, Mahajan-Miklos S, Ausubel FM. 1999. Killing of Caenorhabditis elegans by Pseudomonas aeruginosa used to model mammalian bacterial pathogenesis. Proc Natl Acad Sci U S A. 96(2):715–20. doi:10.1073/pnas.96.2.715.

Trent C, Tsuing N, Horvitz HR. 1983. Egg-laying defective mutants of the nematode Caenorhabditis elegans. Genetics. 104(4):619–647. doi:10.1093/genetics/104.4.619.

Zhou Z, Hartwieg E, Horvitz HR. 2001. CED-1 Is a Transmembrane Receptor that Mediates Cell Corpse Engulfment in C. elegans. Cell. 104(1):43–56. doi:10.1016/S0092-8674(01)00190-8.

